# Prefrontal Manifold Geometry Explains Reaction Time Variability

**DOI:** 10.1101/2023.05.11.540459

**Authors:** Roger Herikstad, Camilo Libedinsky

## Abstract

The stochastic drift-diffusion model proposes that the variability in reaction time is due to randomness during the accumulation of evidence until a decision threshold is reached. However, the neural mechanisms that explain both the randomness and implementation of the decision threshold in the model remain unclear. Here we address these questions using the dynamical systems approach to analyze primate frontal eye field activity and using microstimulation for causal manipulations. We built a mechanistic model in which signals associated with motor plans are bumped out of their attractor state by go-cue signals that emerge ∼60 ms after the go cue. The network then travels through a transition subspace towards a movement-initation subspace that emerges ∼35 ms before movement onset and implements the decision threshold. We postulate that the randomness in evidence accumulation, and hence in reaction times, is explained by the amplification of noise during movement preparation by the geometry of the frontal eye field manifold.

## INTRODUCTION

The length of time it takes for someone to react to a stimulus, or reaction time (RT), varies from trial to trial. Various factors can influence RT variability, such as the level of alertness^1^ and conflicting motor plans^2^. Nevertheless, even when controlling for these factors, there is still significant variation in reaction times^3^. Generally, for well-rested and attentive humans, saccadic eye movements have an RT range between 100 and 350 ms^4^.

The stochastic drift-diffusion model explains the variability in RT across multiple trials as a result of the randomness in accumulating evidence until a decision threshold is reached^5^. Support for this model has been found in multiple brain regions, including in frontal and parietal cortices^6^. For instance, in the frontal eye field (FEF), which plays a key role in eye movement generation, neuron activity increases stochastically towards a specific activity level prior to saccade onset^6,7^. According to the model, the randomness in the rise-to-threshold process accounts for trial-to-trial RT variability^8,9^. Despite its success in capturing many aspects of RT variability^5^ the neural mechanisms that account for trial-to-trial variability in the rise-to-threshold process (and therefore RTs) and the implementation of the decision threshold remain unclear^6^.

To investigate how movements are planned and executed in the brain, we utilized the dynamical systems approach^10–12^. The process of transitioning from planning to executing a movement involves the reorganization of cortical activity, where information encoded in a movement preparation subspace transitions to a near-orthogonal motor execution subspace^13–16^. This near-orthogonality allows for movement preparation without its execution. When sensory cues trigger movements, a non-selective response in motor areas mediates the reorganization of cortical activity^13,16,17^. To identify activity subspaces, it is crucial to find the exact time window when information is encoded within these subspaces. The size of the time window is important since using a window that is too large might blend different activity subspaces. Conversely, a window that is too small may lead to missing relevant information due to noise. Prior studies on movement initiation have typically used relatively large time windows (>200ms) to identify activity subspaces, based on implicit assumptions about cognitive operations and network dynamics^13–16,18^. We hypothesized that a data-driven approach could be used to determine informative time windows, particularly in periods with quickly evolving neural dynamics such as those observed before eye movement execution.

In this study, we investigated the activity of populations of FEF neurons in monkeys performing a delayed-saccade task. Our findings were consistent with previous research, which showed that motor preparation signals are maintained in an attractor state during the delay period^19,20^. Using a data-driven approach, we identified two time windows that were particularly relevant for movement initiation. The first was a 25ms window starting 60ms after the go-cue, which postdicted the go-cue timing. The second was a 35ms window starting 35ms before the movement onset, which predicted the movement onset timing. The variable time between these windows defined a “transition subspace”. Based on these results, we developed a mechanistic model of the neural dynamics and computations involved in movement initiation. In our model, go-cue signals bump motor preparation signals out of their attractor state (∼60 to 85 ms after the go-cue) and the network then drifts through the transition subspace towards a movement-onset state (∼35 to 0 ms before movement), which serves as a threshold for movement initiation.

We hypothesized that minor changes in the network state during motor preparation could be magnified during the transition period. This amplification could occur either by moving along the same trajectory at different speeds (e.g. faster for shorter response times) or by traversing trajectories of different lengths at the same speed (e.g. shorter trajectories for shorter response times), or both. We found that motor preparation signals were indeed predictive of reaction time (RT) as expected^18^, although the amount of explained variance was relatively small. Upon analyzing the length and the speed of the trajectories during the transition period, we discovered that only the trajectory length significantly predicted RT, and it explained several times more variance than the motor preparation activity.

Our model predicts that a slight disruption of the network’s activity during the transition period, but not during the motor preparation period, would result in changes in reaction times of the pre-planned saccades. We tested this prediction by conducting electrical microstimulation of FEF, and we found that quicker pre-planned saccades could be elicited during the transition period, but not before it. These findings provide causal evidence that supports the proposed model.

In summary, our findings give a detailed understanding of how the population dynamics in the FEF contribute to the variability in reaction times and the generation of eye movements. The results suggest that the FEF manifold geometry amplifies the effects of noise fluctuations during the motor preparation period by allowing the network to traverse different paths of varying lengths across trials. Notably, the proposed model is specific to the FEF, as similar results were not observed in the neighboring dorsolateral prefrontal cortex (LPFC), which is consistent with the established role of the FEF in eye movement generation.

## RESULTS

We trained two monkeys (*Macaca Fascicularis*) to perform a visually-guided delayed-saccade task. For one monkey, the delay period contained a distracting stimulus (Monkey J), and for the other monkey the delay period contained either a distracting or a re-target stimulus (Monkey W), although for this study we only analyzed trials in which the second stimulus was a distractor (Figure 1a). The number of correct such trials per session ranged from 86 to 227. The difference between the two tasks is not important because the analysis focused on data collected around eye movements, which was essentially the same for both tasks. Reaction times for both monkeys ranged between 100 ms and 300 ms (Extended data Figure 1). We implanted electrode arrays in the monkeys’ brains in two areas: the FEF, located along the anterior bank of the arcuate sulcus, and the LPFC, located along the dorsal and ventral banks of the principal sulcus (Figure 1b). We analyzed a total of 251 cells recorded from the FEF region,124 from Monkey J and 127 from Monkey W. In addition, we analyzed 202 cells recorded from the LPFC region, 128 from Monkey J and 74 from Monkey W. As previously observed in skeletal motor regions^14,16,17^, activity in the FEF also reorganizes between planning and executing an eye movement, and this reorganization is preceded by a condition-invariant signal (Figure 1c).

**Figure 1.**
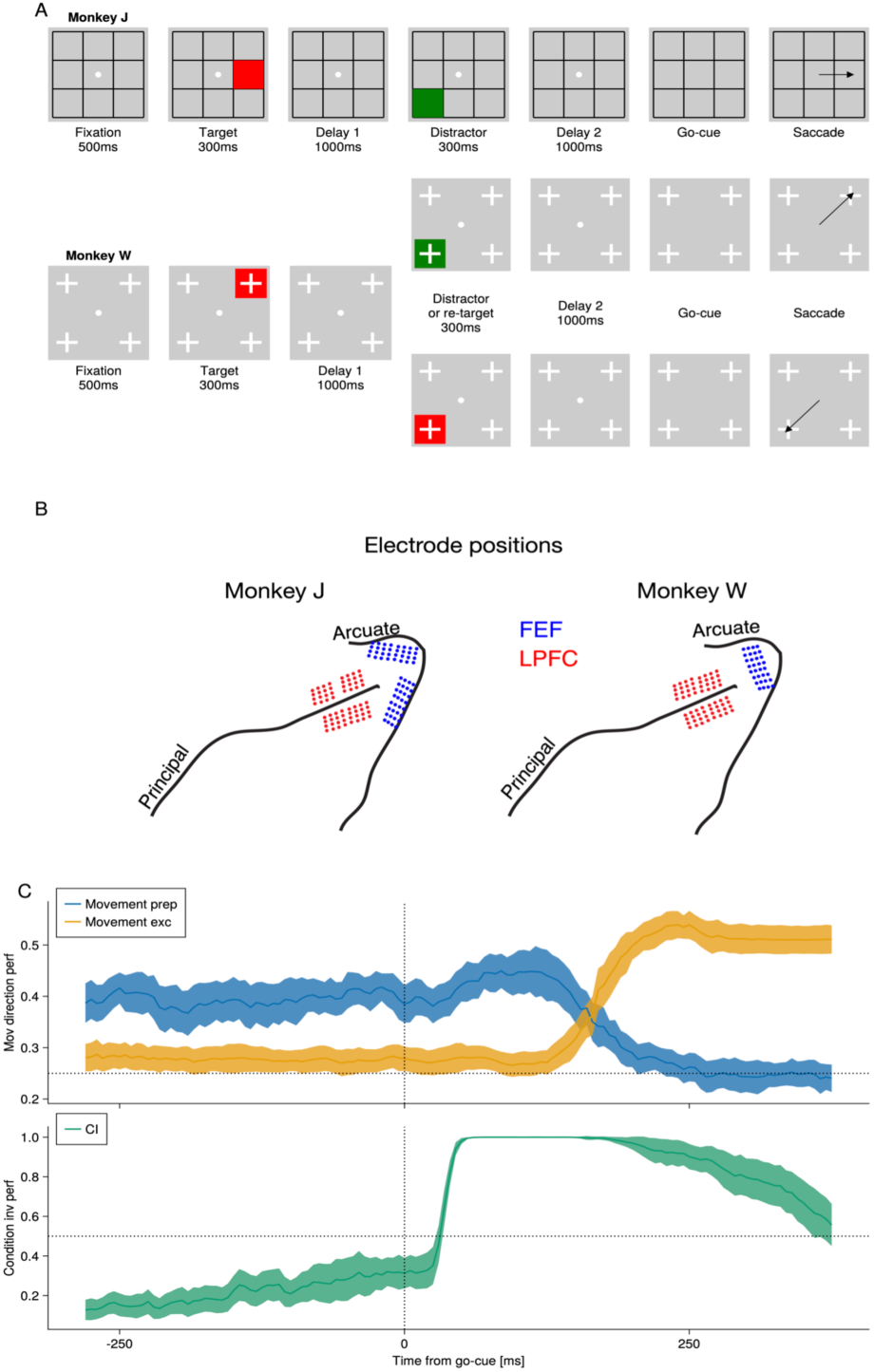
Task Description, electrode locations, and identification of coarse subspaces. **(A)** Task description. Both monkeys performed different versions of a visually-guided delayed saccade task. Trials were initiated by fixating for 500ms on a central fixation spot, after which a red square (or target) was shown for 300ms in 1 out of 8 locations (Monkey J) or 1 out of 4 locations (Monkey W). After a 1000ms Delay 1 period, a second stimulus was presented for 300ms in a different location from the target. For Monkey J, this second stimulus was always a green square, while for Monkey W, the second stimulus had a 50% chance of being a green square and a 50% chance of being a red square. The task rule for both monkeys was to report the location of the last red square seen. Thus, the green square, if shown, served as a distractor. After the second stimulus, a 1000ms Delay 2 period was followed by a go-cue, which was the disappearance of the fixation spot. The monkeys had to saccade to the remembered location within 500ms to receive a juice reward **(B)** Location of implanted electrode arrays. In both monkeys, we chronically implanted electrode arrays in the pre-arcuate region, which includes the FEF (blue), and along the dorsal and ventral banks of the principal sulcus, denoted as DLPFC (red). **(C)** Decoders trained to discriminate different movement plans in the period of −600 ms to 0 ms prior to go-cue (movement preparation), different movement executions in the period of 50 ms to 250 ms after the go-cue (movement execution), and to differentiate activity in the 100 ms before go-cue from the activity in the 100ms after the go-cue (condition-invariant signal). All components were constrained to be orthogonal to each other.

### The dotted lines indicate chance performance

We used a data-driven approach to determine the length and latency of the time windows that were informative about the go-cue and movement onset. Specifically, we looked for time windows in which changes in the activity of the neurons were postdictive of the go-cue timing or predictive of the movement onset timing. To do this, we trained classifiers (using linear discriminant analysis) to distinguish between changes in activity after the go-cue and those during the motor preparation period before the go-cue. We explored a range of window sizes, from 5 ms to 45 ms, and latencies, from 0 ms to 100 ms (Figure 2a for the combined analyses, and Extended data Figure 2 for analyses in single monkeys). For example, a classifier trained to postdict the go-cue onset using a 25 ms window size at a 60 ms latency, was trained to discriminate between the change in population activity between 60 ms and 85 ms post-go-cue and the changes in population activity prior to the go-cue using the same window size. Using this approach, we found that a 25 ms window at a latency of 60 ms post-cue (i.e. from 60 ms to 85 ms after the go-cue), was postdictive of go-cue timing (Figure 2a, left, and Extended data Figure 2). Notice that this is the smallest time window, closest to the go-cue onset. The analysis of single neurons with high contribution to this decoder revealed that some neurons increased, while others decreased their activities during this time window (Figure 2b). Additionally, a 35 ms window at a latency of 0 ms pre-movement (i.e., from −35 ms to 0 ms prior to the movement onset), was predictive of movement timing (Figure 2a, right). Notice that this is the smallest time window, closest to the movement onset. The analysis of single neurons with high contribution to this decoder also revealed that some neurons increase, while others decrease their activities during this time window (Figure 2c). The signals in these two time windows were distinct, meaning that a decoder trained to postdict the go-cue timing could not be used to predict movement onset timing, and vice versa (Extended data Figure 3). The trajectory of the population activity between these two time windows defines a “transition subspace”, and the fraction of explained variance associated with the monkeys’ reaction time peaked during this period (Figure 2d). None of these signals could be identified in the more anterior area DLPFC (Extended data Figure 4).

**Figure 2.**
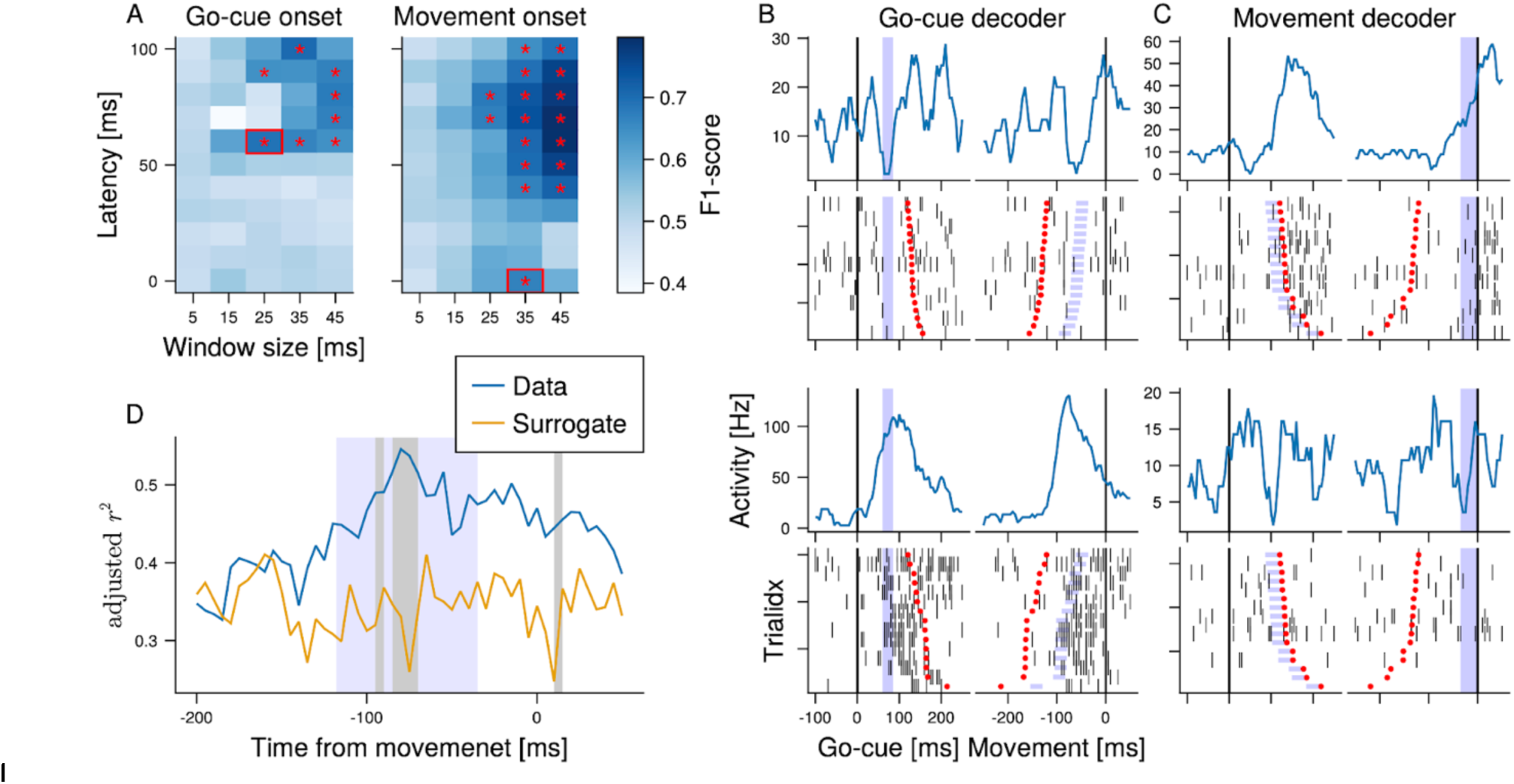
Identification of go-cue and movement-onset signals. **(A)** The F_1_-score of a decoder trained to postdict the go-cue onset (left) and to predict movement onset (right). The red asterisks indicate the combinations of window sizes and latencies where the F_1_-score significantly exceeded 0.5. The bins that are outlined in red correspond to the smallest time windows closest to the event being decoded; in the left plot, it corresponds to the bin closest to the go-cue, and in the right plot it corresponds to the bin closest to the movement onset. **(B)** Examples of cells with high contribution to the go-cue onset decoder during a single session and in a single saccade location, at a window size of 25ms and a latency of 60ms. The purple region highlights the go-cue window (left plots) and pre-movement period (right plots). **(C)** Examples of cells with high contribution to the movement onset decoder at a window size of 35ms and a latency of 0 ms. The first and third rows of (B) and (C) show the PSTH aligned to go-cue onset (left) and movement onset (right), while the second and fourth rows show the rasters for the same alignments. The blue shaded area represents the window used to train and test the decoder, while the red dots indicate either the movement onset (left) or the go-cue onset (right). **(D)** The mean adjusted r^2^ for data (blue) and surrogates (orange). Adjusted r^2^ was computed separately for each session and location. The gray shaded areas indicate windows in which the distribution of adjusted r^2^ for the data was significantly greater than that of the surrogates (rank sum test, p < 0.01). The blue shaded area represents the extent of the transition period from go-cue space to pre-movement space.

**Figure 3.**
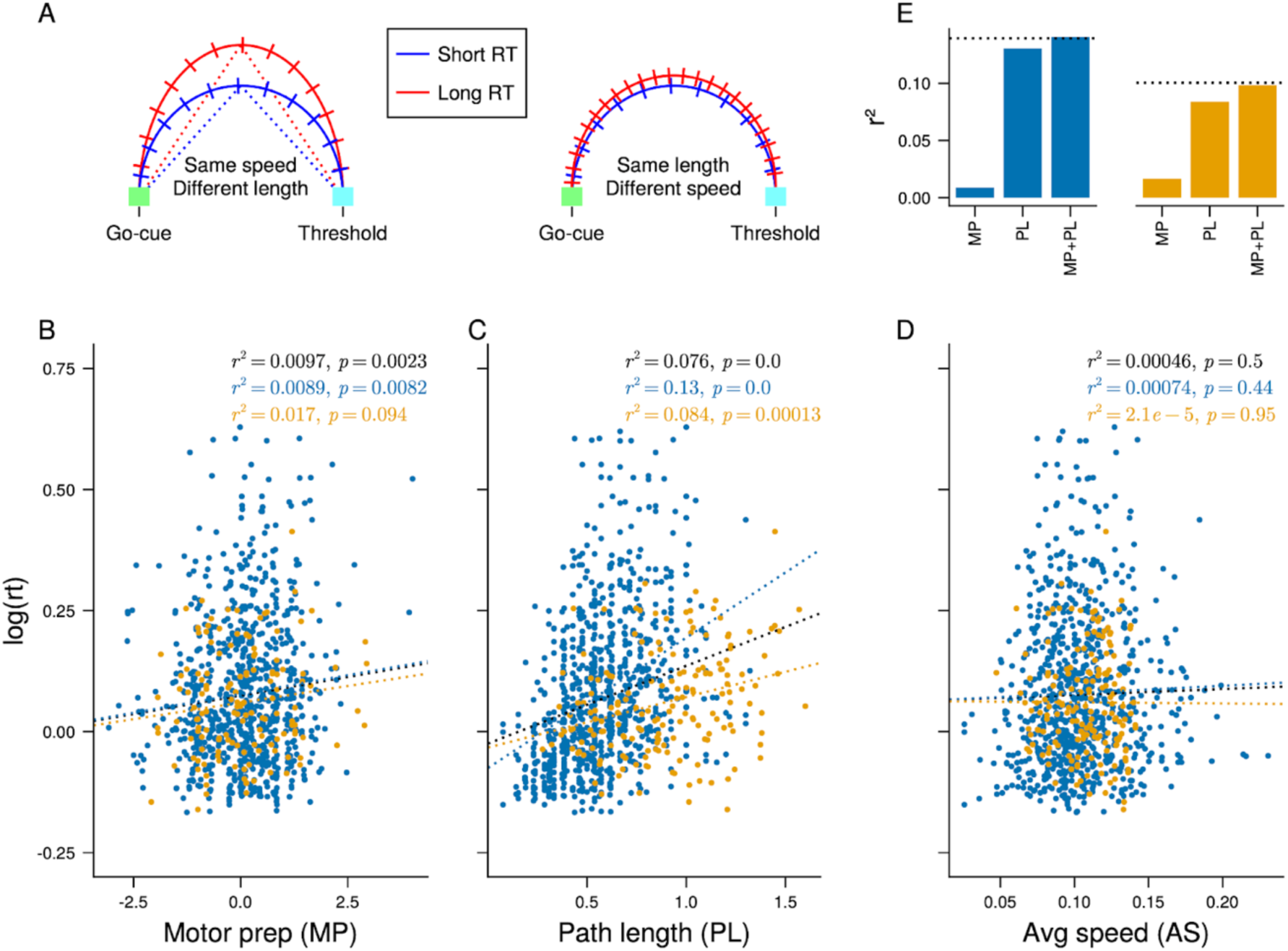
Prediction of reaction times. **(A)** Illustration of two possible ways to explain reaction time variability using population trajectories; one associated with short reaction time (blue) and one with long reaction time (red). The curved lines represent population trajectories, and the ticks along the trajectories demark time. Left: The long reaction time results from a longer trajectory at a constant speed, compared to the short trial. The dotted lines illustrate how path length was computed, by finding the point on each trajectory that maximized the sum of the Euclidean distance from that point to the start and end point of the trajectory. Right: The longer reaction time trial results from the slower speed of the same trajectory (i.e. more elapsed time) compared to the short reaction time trial (the lines do not overlap for illustration purposes only). **(B-D)** Regression to reaction time using movement preparation activity (projection onto 1st factor) (B), transition period path length (C), and transition period path speed (D), across all subjects, sessions, and locations. Monkey W in blue, monkey J in orange, and both monkeys combined in black. **(E)** r^2^ for regression using both movement preparation (MP) activity and path length (MP + PL), compared to regression using either variable on their own (MP and PL). The dotted horizontal lines indicate the expected r^2^ value if the two variables were independent.

**Figure 4.**
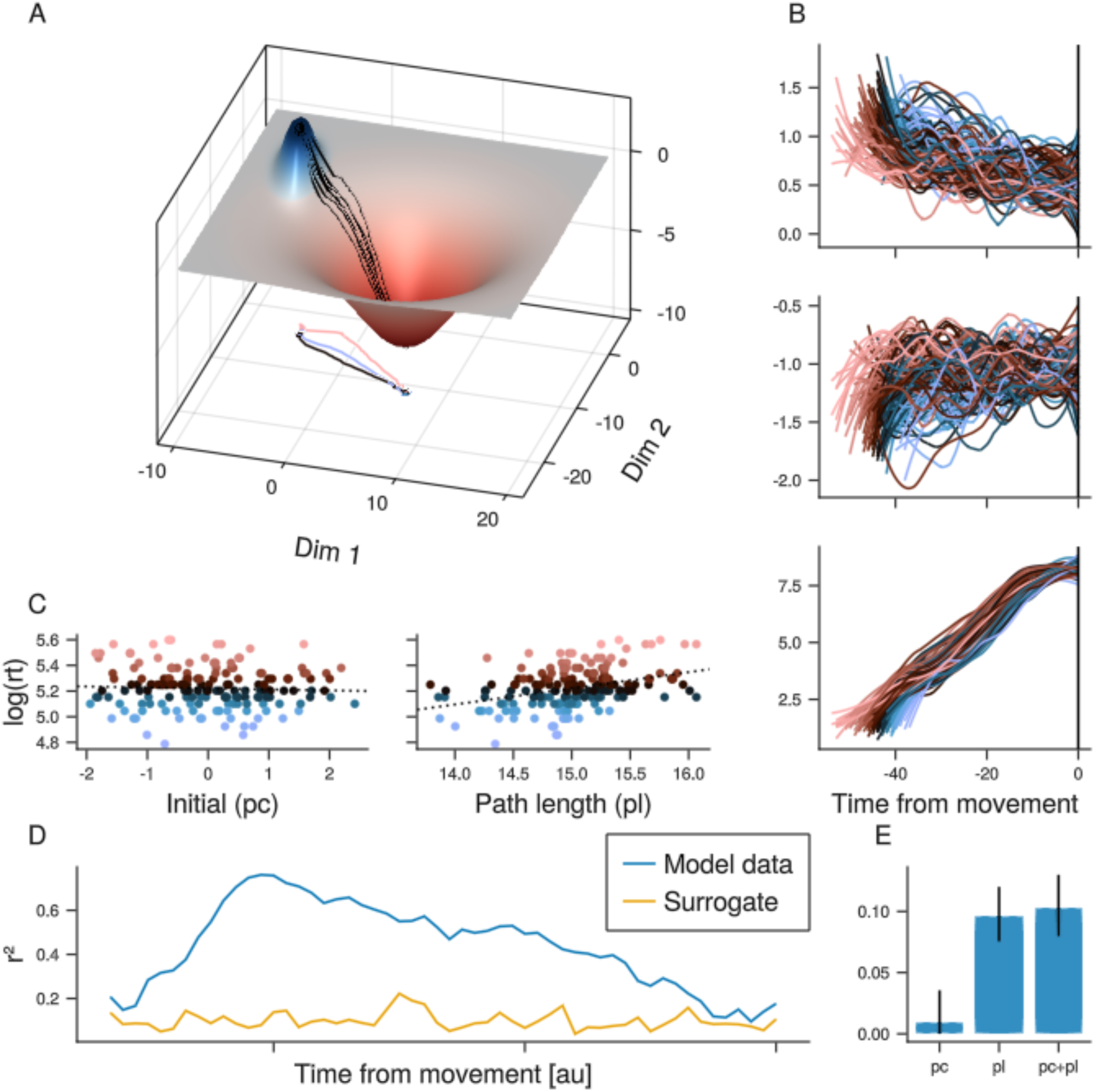
Network Model. **(A)** The 3D manifold constraining the population responses as they move from the delay 2 basin (left, blue) to the pre-movement attractor (right, red). Each black line is the trajectory followed in one trial. **(B)** Examples of trial-by-trial responses of 3 cells. These responses were obtained by projecting the 2D trajectories shown in (A) onto a 15-dimensional space via an orthonormal projection matrix, then adding gaussian noise. Each line in these plots represents the smoothed activity for one trial. **(C)** The logarithm of reaction time is plotted against initial conditions (left) and path length (right) of the activity in the 15-dimensional space. A one-dimensional representation of the initial conditions was obtained by projecting each trial onto the space spanned by the first factor in a Factor Analysis. **(D)** Fraction of explained reaction time variance (r²) as a function of time using the full, i.e., 15-dimensional response at each time point. The orange trace shows the r² value when the reaction time was shuffled across trials. **(E)** r^2^ for regression using both post-cue activity and path length (pc+pl), compared to regression using either variable on their own (pc and pl) for 50 iterations (error bar denotes the 5th and the 95th percentile of a distribution obtained by projecting onto higher dimensional spaces and adding noise consistent with the number cells and trials in the neural population data over 500 runs).

To investigate whether FEF activity could predict reaction time, we examined the activity during two different periods: the motor preparation period (0 ms to 50 ms after the go-cue) and the transition subspace period. During the motor preparation period, we performed factor analysis (FA) and found that one dimension commonly accounted for over 90% of the variance associated with reaction time^18^ (Extended data Figure 5). We then used a projection of population activity onto the first factor to carry out a regression analysis and found a small but significant correlation with reaction time (r^2^: Monkey W: 0.0089, p<0.01; Monkey J: 0.017, n.s. p<0.1; both monkeys combined: 0.0097, p<0.01; Figure 3b). For the transition period, we analyzed the speed and length of the trajectories (Figure 3a). We found that only trajectory length was correlated with RT (r^2^: Monkey W: 0.13, p<0.01; Monkey J: 0.084, p<0.01; both monkeys combined: 0.076, p<0.01; Figure 3c,d). Multiple regression of both motor preparation and transition period length revealed that these two factors were mostly independent (Figure 3e).

**Figure 5.**
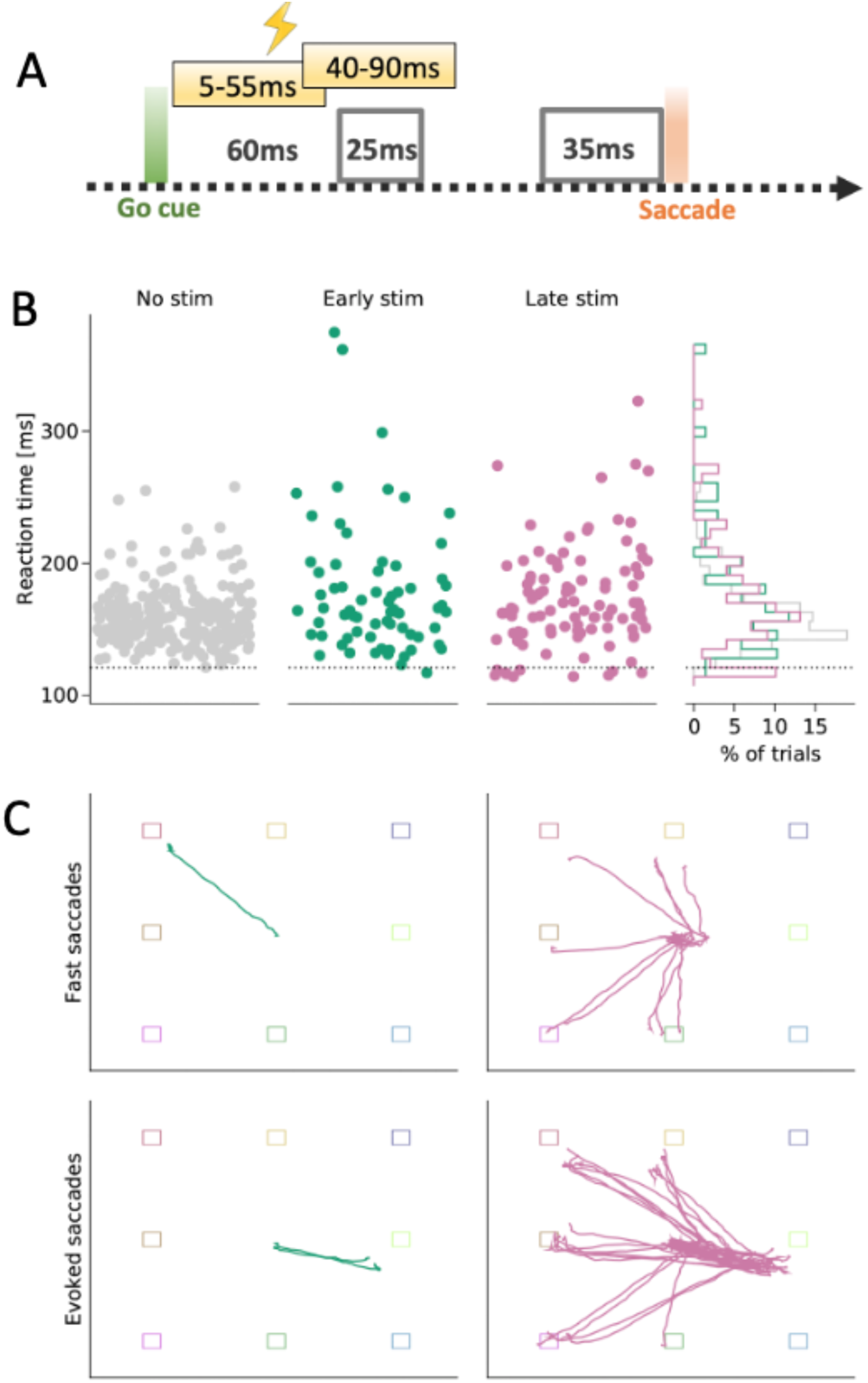
Microstimulation. **(A)** A 12-40 mA, 50 Hz, 50 ms stimulation pulse was delivered simultaneously to all 32 electrodes either starting at 5 ms or 40 ms post go-cue onset. Gray boxes indicate the timing of the go-cue signals (left) and movement-onset signals (right). **(B)** The scatter plots show the reaction times for non-stimulated trials (light gray), trials where the stimulation started at 5 ms (green), and at 40 ms (purple), for trials in which the monkey executed a saccade to the correct target location. The histogram shows that the reaction times for the stimulated trials exhibited a bimodal distribution, with a group of trials associated with a shorter reaction time compared to the non-stimulated trials (<121ms; which is the minimum of the no-stimulation reaction time distribution, represented by the dotted horizontal line). **(C)** The details of these short-latency saccades to the correct location are shown in the bottom panels, where the eye trajectory during each saccade is overlaid on a display of the possible target locations. Green traces correspond to trials with early stimulation (starting 5 ms after go-cue), while purple traces correspond to trials with later stimulation (starting 40 ms after the go-cue). The top panels show all the fast saccades (below the dotted line in B), that went directly from fixation to target location. The bottom panels show all the evoked saccades triggered by stimulation, as well as the subsequent saccades.

### Data from monkey W in blue, from monkey J in orange

A dynamical system model integrates these results to explain the mechanisms of eye-movement initiation and RT variability (Figure 4). During motor preparation, the network’s activity is maintained in an attractor state. The go-cue signal bumps the activity out of this attractor state around ∼60 ms after the go-cue. The network state then travels through the transition subspace towards the pre-movement subspace around ∼35 ms prior to movement, which serves as a threshold for movement initiation. The variability in reaction time is caused by the network taking different trajectories in the transition subspace, such that longer trajectories lead to slower RTs. These different trajectories are caused by small noise fluctuations in the motor preparation attractor state. The results shown in Figure 3e, which show that contributions of motor preparation signals and the trajectory length during the transition period were mostly independent, appear to be at odds with our model, in which transition period trajectory lengths are determined by the initial conditions during motor preparation. However, analysis of the model confirmed that the observed result is indeed consistent with the model, given the noise levels found in the data (Figure 4b-e).

Our proposed model suggests that perturbing the network using electrical microstimulation during the transition period would alter the reaction times of the monkey, possibly making them faster. On the other hand, perturbing the network during the more stable movement preparation attractor state would have minor behavioral effects. To test these predictions, we performed large-scale low-current microstimulation of the FEF, injecting current in all 32 FEF electrodes simultaneously. We found that precisely timed brief pulses could trigger pre-planned eye movements that were faster than normal, but only when the stimulation occurred during the transition subspace period and not earlier, which is consistent with the proposed model (Figure 5).

## DISCUSSION

Explaining how abstract psychological models are instantiated by particular neural circuits has been a longstanding goal of cognitive neuroscience. One such model is the stochastic drift-diffusion model, which has successfully explained different aspects of decision-making and movement initiation^6^. This model involves abstract concepts like the stochastic accumulation of evidence and a decision threshold. Previous studies have shown that some neurons in the FEF exhibit behaviors that are consistent with the stochastic drift-diffusion model, such as an increase in activity before movements (indicative of the stochastic accumulation of evidence), and reaching a certain activity level before the movement (indicative of a decision threshold^7^). However, to date, we have lacked a mechanism to explain how these abstract objects are implemented in neural networks.

Here we addressed this gap by developing and testing predictions of a mechanistic model that explains how eye movements are initiated, and why they vary from trial to trial. We used neurophysiological recordings of FEF neurons and a dynamical systems analysis of their responses to build the model. We propose that the variability of the time taken to initiate an eye movement is caused by the amplification of neural noise by the geometry of the FEF manifold. Specifically, we propose that reaction time variability is the result of the different trajectory lengths of population activity through a transition subspace, defined as the activity space between go-cue signals (∼60 to 85ms after the cue) and a movement-initiation subspace (∼35 to 0ms before the movement onset). The timings of these different time windows are consistent with previous behavioral and neural observations. For instance, the movement-initiation subspace, which would implement the decision threshold in the stochastic drift-diffusion model, occurs in the period between −35 ms to 0 ms before movement, which is consistent with the expected time window in which FEF can influence eye movements^7,21^. In addition, the transition subspace onset, which marks the point-of-no-return for saccade initiation, starts around 100 ms before the saccade, which is consistent with the observation that planned eye movements can be canceled up until 100 ms before the saccade, but not later^22,23^.

Under the dynamical systems approach, a network can be characterized by its initial conditions, its inputs, and rules, or equations that determine how the network’s state changes over time^11^. In our model, the initial conditions correspond to the movement preparation state prior to the go-cue signals, the inputs correspond to the go-cue signals, and the rules that govern the evolution of the network’s state over time are reflected in the network’s manifold and dynamical landscape. Previous studies have shown that movement preparation signals are predictive of reaction time in the FEF^18^, motor^24–26^, premotor, and parietal cortex^27^. We were able to confirm these findings in the FEF (Figure 3). We also found that the trajectory length during the transition period was correlated with reaction time and that the variance explained by the transition period trajectory length was several times larger than the variance explained by the movement preparation period. This led us to hypothesize that small noise fluctuations during the movement preparation period (or initial conditions) are amplified by the geometry of the FEF manifold, and this amplification is reflected in longer trajectory lengths. In a different prefrontal region, the dorsomedial frontal cortex (DMFC), initial conditions in conjunction with context-dependent tonic inputs dictate the speed, but not the trajectory length, of neural dynamics for reproducing timing intervals^28^. This difference may reflect different strategies that can be used by neural networks to adjust the temporal patterns of neural dynamics. One strategy may rely solely on network-intrinsic properties (the network’s manifold and dynamical landscape), while another strategy may rely on both network-intrinsic properties and network-extrinsic inputs, to control the temporal patterns of neural dynamics. We speculate that context-dependent modulations of reaction time could be driven by a combination of both strategies. For instance, highly-motivated subjects may respond faster than less-motivated ones^29–31^. This urgency could be reflected in a tonic input to the FEF that affects the speed of the trajectories in the transition subspace, while at the same time maintaining the variability across trials due to the geometry of its manifold.

The dynamical systems approach also allowed us to address the question of how FEF neurons can perform different functions at different stages of a task^6^. For example, some neurons may initially be involved in evidence accumulation during movement planning and then, subsequently, in movement execution^32^. This presents a paradox if we consider single neurons independently because it’s unclear how neurons can shift their roles during a task. However, our dynamical systems model of the FEF network provides an explanation for this apparent paradox. We show that the changes in single neuron selectivities over time can be explained by the orderly transition of the network state across different activity subspaces. This means that the function of an individual cell is interpreted in the context of the activity of other neurons at different stages of the task, rather than in isolation.

The group of neurons that control eye movements are distributed across several brain regions, including cortical regions in the prefrontal and parietal cortex, and subcortical regions like the superior colliculus, thalamus, and brainstem oculomotor nuclei. In the present study, we only measured the activity of FEF neurons, but we interpret our analyses in the context of the broader network involved in eye movement control. For instance, motor preparation signals in FEF may involve an interaction with other brain regions, such as the thalamus^33,34^, or the DLPFC^35^. Similarly, we propose that the movement-initiation subspace might function as a decision threshold, but it would only serve as such if downstream regions, like the superior colliculus, selectively extract information from this subspace^12^. This means that the subspace we discovered is a putative threshold, and conducting simultaneous measurements with downstream regions would be needed to confirm it as such.

To test the model, we investigated whether stimulation during the transition period could impact the speed of responses by using low-current microstimulation of FEF electrodes. We found that stimulation during the transition period, but not the preceding motor preparation period, led to faster initiation of pre-planned movements in a subset of trials. Interestingly, saccades that were sped up were always directed towards the hemifield ipsilateral to the stimulation site in the left hemisphere (Figure 5). In other trials, saccades were evoked by the stimulation towards a fixed location in the contralateral (right) hemifield. After this stimulation-evoked saccade, the monkeys made a corrective saccade toward the remembered original target location. Like the pre-planned saccades, these corrective saccades were also directed towards the hemifield ipsilateral to the stimulation site in the left hemisphere. Interhemispheric callosal connections between the left and right FEFs are inhibitory for conflicting saccades^36^. Therefore, inputs from the contralateral FEF could be viewed as a contextual signal carrying information about the preparation of ipsilateral eye movements. Microstimulation of the contralateral hemisphere could disrupt this contextual signal and affect the speed of the neural dynamics^28^, which in turn could impact the reaction time. The corrective saccades observed after the stimulation-evoked saccades were likely driven by the saccade plan encoded in the contralateral hemisphere, which was then executed after the stimulated hemisphere triggered the evoked saccade. However, contralateral corrective saccades could not be executed since the stimulation likely interfered with the motor plan toward those targets.

Our results raise several questions. For example, are the mechanisms we have identified for initiating eye movements also applicable to the initiation of skeletal movements? In tasks that require the gradual accumulation of evidence before initiating a response^37^, does the decision threshold correspond to the entry into the transition subspace, or into the movement-initiation subspace? Moreover, can we view the go-cue signal as a fast and unambiguous type of evidence accumulation, while tasks that involve slow evidence accumulation simply progress slowly towards the transition period? The go-cue signal that we described is the result of a change in the sensory stimulus, specifically the disappearance of the fixation spot. However, this signal may actually represent a decision made by the animal to initiate the saccade rather than being solely a sensory response. This is because the animals can choose to ignore the go-cue if they desire. If this is true, then this decision signal should also be present in self-initiated saccades. Further research will be required to answer these intriguing questions.

## METHODS

### Subjects and surgical procedures

We used two adult male macaques (*Macaca fascicularis)* in this experiment; Monkey J (age 4) and Monkey W (age 12). All animal procedures were approved by, and conducted in compliance with, the standard of the Agri-Food and Veterinary Authority of Singapore and Singapore Health Services Institutional Animal Care and Use Committee (SingHealth IACUC #2012/SHS/757), and the National University of Singapore Institutional Animal Care and Use Committee (NUS IACUC #R18-0295). Procedures also conformed to the recommendations described in *Guidelines for the Care and Use of Mammals in Neuroscience and Behavioral Research* (National Academies Press, 2003). Each animal was first implanted with a titanium head-post (Crist Instruments, MD, USA) before arrays of intracortical microelectrodes (MicroProbes, MD, USA) were implanted in multiple regions of the left frontal cortex. In Monkey W, one array of 32 electrodes was placed over the FEF, and two arrays of 32 electrodes each were placed over the DLPFC, while in Monkey J, 2 arrays of 32 electrodes each were placed over the dorsal and ventral aspect of the FEF, and 2 arrays of 32 electrodes each were placed over the DLPFC (as shown in Figure 1a). The arrays consisted of platinum-iridium wires with either 200- or 400µm separation, 1-5.5 mm long and with 0.5MΩ of impedance, arranged in 4 x 4 or 8 x 4 grids.

### Recording techniques

Neural signals were initially acquired using a 256-channel Plexon OmniPlex system (Plexon INc., TX, USA) with a sampling rate of 40kHz (Monkey J), or a 128 channel Grapevine recording system (Ripple) with a sampling rate of 30kHz (Monkey W). The wideband signals were bandpass-filtered between 300 and 3,000 Hz, and spikes were detected on each channel separately using an automated sorting algorithm based on Hidden Markov modeling^38^. Eye positions were obtained using an infrared-based eye-tracking device from SR Research Ltd (Eyelink 1000 Plus for monkey J and Eyelink 2000 for monkey W). The behavioral task was controlled by a standalone PC, using either the Psychophysics Toolbox in Matlab (Mathworks, MA, USA) (Monkey J), or PsychoPy in Python^39^ (Monkey W). To align the neural and behavioral activity (trial epochs and eye data) for data analysis, we generated digital markers denoting epochs and behavioral responses (break fixation, correct or incorrect response) during the trial. These markers were sent to the Plexon/Grapevine systems using a parallel port, and to the Eyelink system using an ethernet cable.

### Microstimulation

For arrays positioned in the pre-arcuate region (FEF), we used standard electrical microstimulation to confirm that saccades could be elicited with low currents in monkey J (however, we could not perform this procedure in monkey W). The procedures and parameters used have been reported previously^40^.

We also used electrical microstimulation in a third animal, Monkey Z, for the results shown in Figure 5. This animal was trained on the same task as Monkey W. Specifically, we used bi-phasic pulses consisting of 200µs of negative phase, followed by 100µs of 0 phase, and 200µs of positive phase, at an amplitude of 12 to 40 µA. These pulses were delivered in short bursts of 50ms at a frequency of 50Hz, at 20 ms, 5 ms, or 40 ms after the presentation of the go-cue, across all 32 stimulation channels.

### Behavioral tasks

For Monkey J, targets and distractors were presented on a 3 x 3 grid. After a 500ms pre-fixation period, a red square, constituting the target, was flashed for 300ms in one of the 8 outer positions on the grid. A 1s delay period followed, after which a green square, constituting the distractor, was flashed for 300ms in one of 7 positions (target and distractor never occupied the same position). Then, another 1s delay period followed, before the disappearance of the central fixation spot (the go-cue) indicated to the animal that he should respond with an eye movement to the remembered target position. To receive the reward, constituting a drop of fruit juice, the animal had to maintain fixation on the target position for 200ms (Figure 1a top).

The trial structure for Monkey W was similar, but with 3 key differences. First, the animal was presented with crosses at the target positions instead of a grid outline indicating the possible target positions. Second, the animal was only presented with 4 possible target positions, instead of the 8 presented to Monkey J. Third, the second stimulus could either be a new target (50% chance), or a distractor (50% chance), instead of only distractors (Figure 1a bottom).

When comparing results across the two animals, we combined pairs of target positions for Monkey J to create 4 groups of 2 target locations each.

### Orthogonal subspaces

To find mutually orthogonal subspaces for delay 2 and pre-movement target location information, as well as information about whether or not the go-cue had been presented (Figure 1C), we followed the approach in Cunningham & Ghabramani (2015)^41^ to train an orthogonalized multi-class linear discriminant decoder. To that end, following the standard multiclass LDA approach, we first constructed between-and within-class scatter matrices for target location using the last 600ms of the delay 2 (d2) period, as well as the first 400ms of the post-go-cue (pm) period. In addition to this, we also constructed scatter matrices for the difference between activity in the last 100ms of the d2 period, and the first 100ms after the go-cue onset. We then ran an optimization to find the projection matrices *W_d2_*. *W_pm_*, and *W_gc_* that jointly maximized:

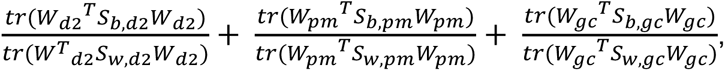

subject to *W^T^_d2_ W_d2_*= 1, *W^T^_pm_ W_pm_*= 1, *W^T^_gc_ W_gc_*= 1, *W^T^_d2_ W_gc_*= 0, *W^T^_d2_ W_pm_*= 0, *W^T^_gc_ W_pm_*= 0. Here, *S_b,x_* is the between-class scatter matrix for the period *x* (i.e. d2 = delay2, pm = pre-movement, and gc = go-cue), while *S_w,x_* denotes the within-scatter matrix. The scatter matrices within each period were constructed using the activity of a pseudo-population where trials were matched either by target location (for d2 and pm), or by go-cue onset (for gc).

### Identification of Go-Cue and Movement onset signals

In the results shown in Figure 2a we sought to identify the time window in which the population activity was maximally postdictive of the timing of the go-cue and another window that was maximally predictive of when the upcoming eye movement was going to occur. To identify points in time where the neural population response was predictive or postdictive of certain events, e.g. eye movement or go-cue onset, we looked for windows in which the population responses were maximally predictive (or postdictive) of those events. Since we were specifically looking for onset signals, rather than using the population responses themselves, we focused on the slope of the responses. Then, for a specific window size and latency, we trained a linear discriminant decoder to separate the slope at that latency from the slope at a random window (using the same window size) in the delay 2 period. In this way, the decoder was constructed to identify changes in population activity aligned to specific events.

To quantify the performance of the decoder, we used the *F_1_*-score, given by

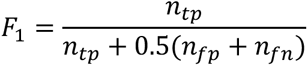

where *n*_tp_ is the number of true positives, *n*_fp_ the number of false positives and *n*_fn_ the number of false negatives. For a decoder that is simply guessing, the number of true positives, false positives and false negatives is the same, and so the *F*_1_ score for such a decoder is 0.5. In our case, a true positive occurred when the decoder correctly concluded that a given response slope originated at a specific latency relative to an event (go-cue onset or eye movement), a false positive occurred when the decoder incorrectly concluded that a slope from the baseline originated at a specific latency relative to an event, and a false negative occurred when the decoder incorrectly concluded that a slope occurring at a specific event latency originated from the baseline.

### Characterizing the transition period

We tested whether the neural activity during the transition period from movement planning to execution carried information about reaction time. First, we computed the average speed of the neural trajectory by summing up the displacements from one point of the trajectory to the next, then dividing by the number of points. For path length, we used a procedure that eliminated the bias caused by the fact that trajectories associated with longer reaction times necessarily had more points, and therefore path length computed as the sum of displacements was trivially correlated with reaction time. Instead, we reduced each trajectory to 3 points; the first and the last point were identical to the actual first and last point of the trajectory, while the third point was identified by maximizing the sum of the Euclidean distances between that point and the first and the last points (Figure 3a).

### Correlation between reaction time and neural responses

To establish whether neural responses contained information about the timing of the saccade relative to go-cue onset, i.e. reaction time of the animal, we performed a linear regression of reaction time *t_r_* to neural responses.

To establish the time course of correlation between neural responses and reaction time (Figure 2d), first, we aligned the neural responses to movement onset, and computed spike counts in 20ms windows from 200ms before movement onset, using steps of 5ms. For each window, we then computed coefficient vectors *β* such that

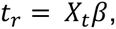

where *t_r_* is the reaction time, *X_t_* is a *n_trials_* x *n_c_* matrix representing the spike counts of *n_c_* cells at time point *t* across *n_trials_* trials. We estimated *β* for each session and each location. To avoid overfitting due to a limited number of trials, we used ridge regression to find *β*, with a fixed ridge coefficient λ of 0.05. We used a modified degrees-of-freedom adjusted r^2^ to determine the goodness-of-fit, where the degrees of freedom *p* was given by^42^:

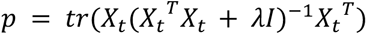

We also examined the relationship between reaction time and neural responses in specific windows (Figure 3b-d). First, we used a 50ms window starting at the onset of the go-cue and counted the total number of spikes in this window across all trials. This served as our approximation of the initial conditions for the transition from delay 2 to movement generation. Prior to running the regression of reaction time, we reduced the dimensionality of these responses using Factor Analysis^18^. Using a two-step process to determine the number of factors^18^, we found that for most sessions, 90% of the shared variance in the post-cue responses could be explained using a single factor Extended data Figure 6). The projection of the neural responses onto this factor was then used to regress the reaction time. To ensure that the relationship between the factor projection and reaction time was consistent across sessions, we estimated the slope between factor projection and reaction time for each session and location. If the slope was negative, we flipped the sign of the projections prior to including them in the overall regression. Factor projections, path lengths, and reaction times were then collated across sessions and locations, and the regression analyses were performed on these collated datasets.

Finally, we used a nested regression approach to test whether the information about reaction time contained in the initial conditions was also contained in the path length. That is, we compared the residual of a regression model consisting of both initial conditions and path length to one containing only path length. If path length contained all the information about reaction time, including the information represented by the initial conditions, the decrease in residual from the combined model would not be greater than that expected by chance, simply by adding more variables. We used the F-test to compare the residuals, where the quantity of interest, *F*, is given by:

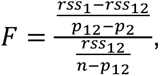

where *rss_1_* is the residual from regression using only path length, *rss_12_* is the residual of the combined regression model, *p_1_* and *p_12_* are the degrees of freedom of the two models, and *n* is the number of observations.

### Model

As a conceptual illustration of how the transition from the Delay 2 period to motor execution might happen, we focused on a toy model consisting of two attractor states. The Delay 2 attractor state was described by a difference of two Gaussian bumps. This provided a stable state in which the memory of the target location resided. The movement onset attractor state was modeled as an inverted Gaussian, with a basin that was lower than the delay 2 basin. The combined shape of these two basins constituted the neural manifold along which the activity of the two model neurons was constrained to evolve. We then postulate that the appearance of the go-cue, in the form of the central fixation point vanishing, temporarily increased the activity of the delay 2 state, enabling it to escape from the delay 2 attractor basin and “roll” down the hill towards the movement onset attractor basin. We further assumed that the exact position in state space prior to the response cue onset was subject to noise, resulting in a variability in the direction at which the neural state escaped the basin. For those trials in which the neural state escaped the delay 2 basin close to the movement onset basin, the path was shorter than for those trials in which the neural state was further away. For these latter trials, the neural state traveled further along the manifold, thus resulting in a longer reaction time, than for the former trials.

In addition, we made a few key modifications to the Delay 2 basin to replicate the data findings as closely as possible. Notably, with a pure difference-of-gaussian delay 2 basin, the neural state would accumulate near the bottom of the basin and be distributed symmetrically around the center. In the data, we found that the initial conditions projected onto a 1-dimensional space (i.e. the first factor), contained a significant amount of information about reaction time. To replicate this in the model, we first made the bottom of the basin flat. This ensured a larger spread in the initial conditions. Secondly, we made the shape of the basin elliptical rather than circular (for the manifold shown in Figure 5, the eccentricity was 2), which introduced an asymmetry that the factor analysis could exploit. Lastly, we oriented the major axis of the ellipse at an angle with respect to the line connecting the center of the delay 2 basin to the center of the pre-movement attractor (for the manifold shown in Figure 5, the angle was 135 degrees). This ensured asymmetry in the path lengths, i.e. that trajectories that originated equidistantly from the line connecting the delay 2 basin and the pre-movement attractor did not result in identical path lengths.

All analyses were implemented in the Julia programming language^43^.

## ACKNOWLEDGEMENTS

We acknowledge Shih-Cheng Yen, Weicheng Tao, and Sherry Aw for the discussions and useful comments on the manuscript.

## EXTENDED DATA

**Extended data Fig 1.**
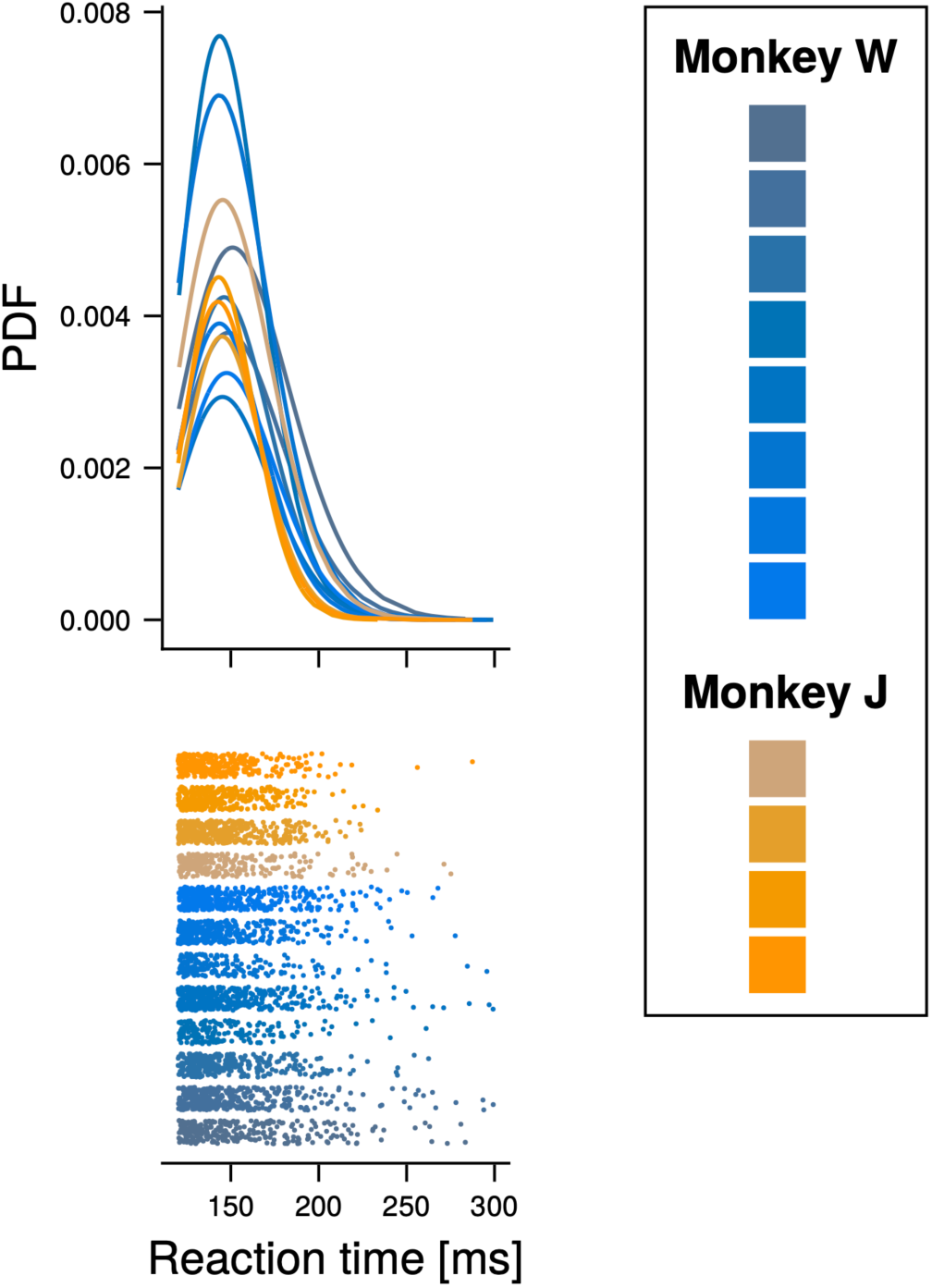
Reaction time distribution. **(Top)**. Distribution of reaction times (probability density function: PDF) for individual sessions of monkey W (blue) and monkey J (orange). **(Bottom)** Reaction times for the distributions shown above.

**Supp Fig 2.**
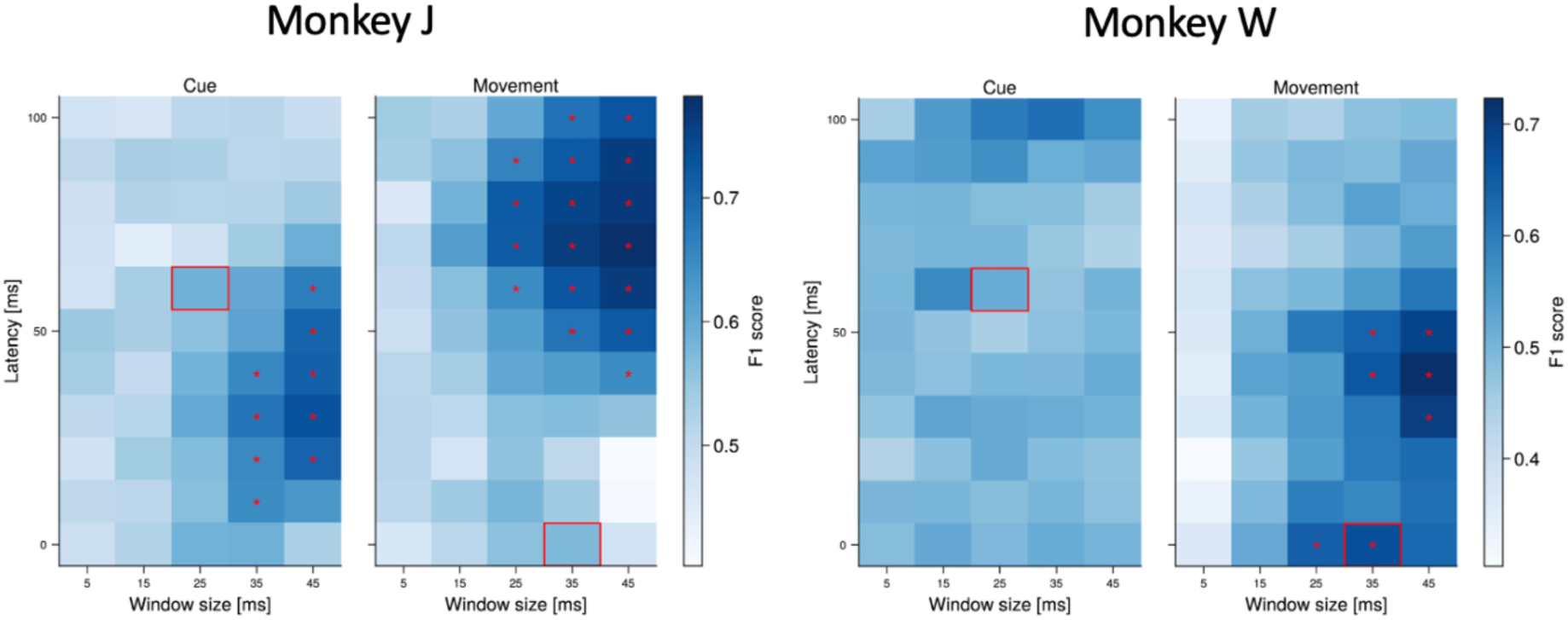
Single Monkey Decoding plots. The same analysis of Figure 2a is shown here for each monkey separately, for cue-aligned (left plots) and movement-aligned (right plots) decoding. The red square highlights the latency and window size combinations identified in the combined analysis in Figure 2a.

**Extended data Fig 3.**
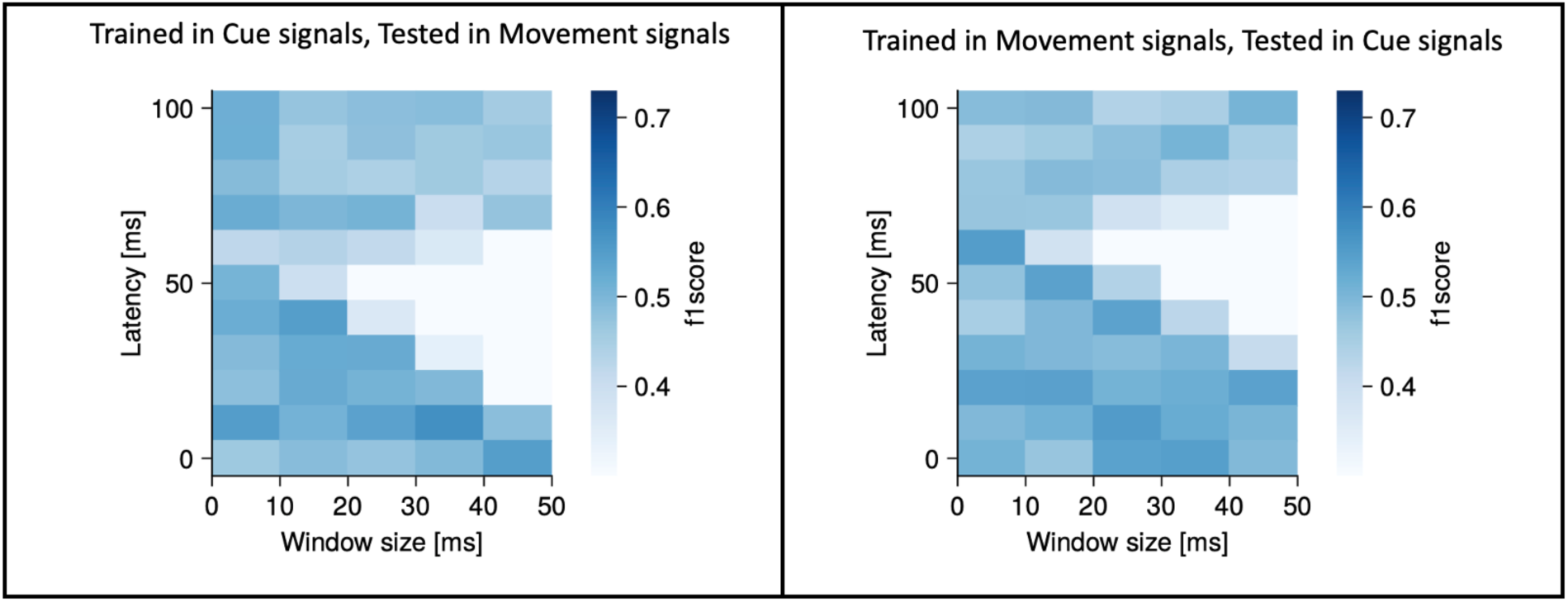
Cross-period decoder. Performance of the same decoders shown in Figure 2a, but with different training and testing periods. The left plot was trained on cue-period signals and tested in movement-period signals (as identified in Figure 2a), and the right plot does the reverse. No values were significantly above 0.5, indicating that the decoder performed at chance level. This figure shows that the signal changes that are postdictive of the go-cue timing are not the same as those that are predictive of movement onset.

**Extended data Fig 4.**
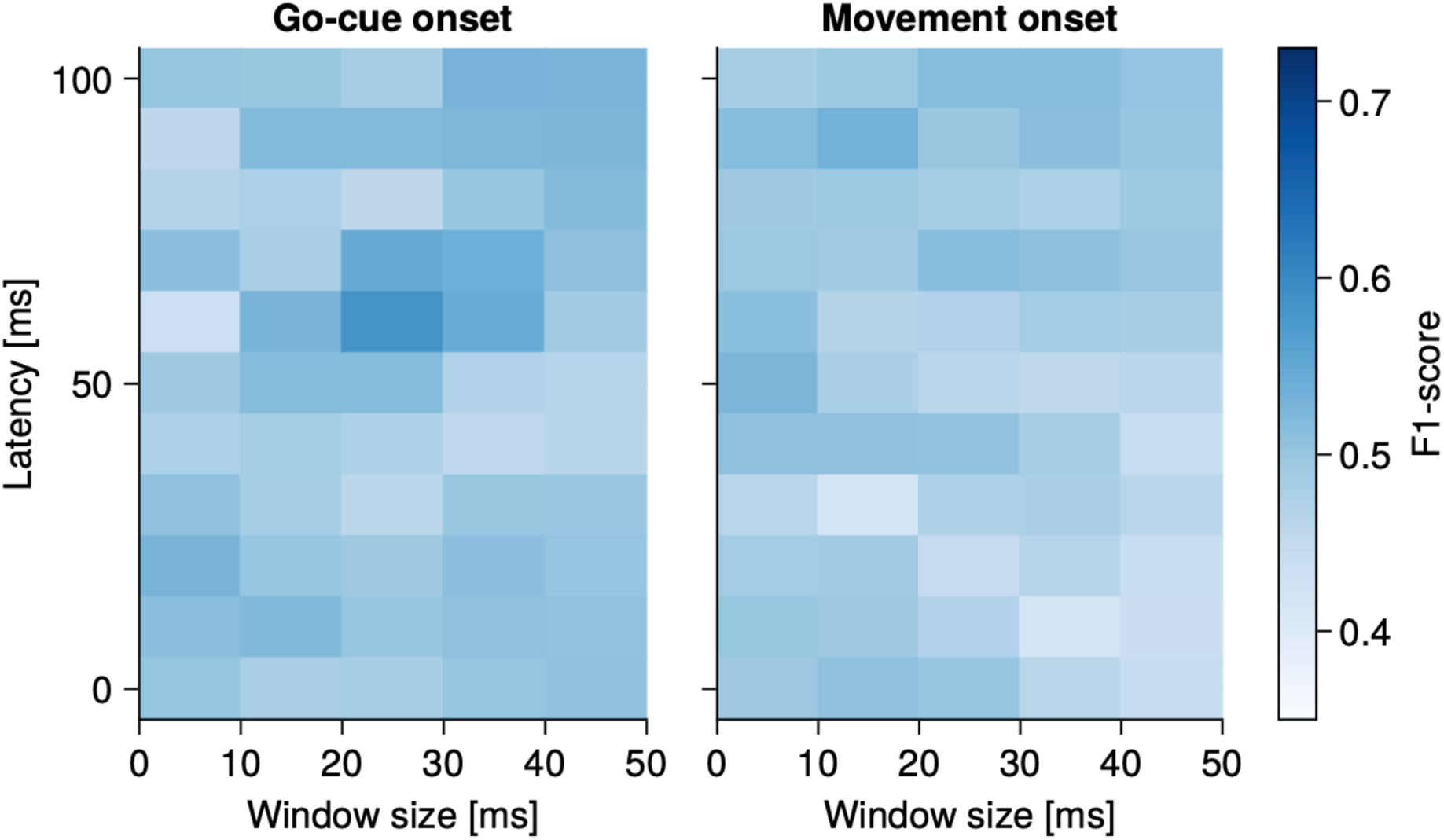
DLPFC does not have go-cue nor movement onset signals. The same analysis that was run in Figure 2a in the FEF was carried out in the DLPFC. No values were significant, so these periods cannot be identified in the DLPFC.

**Extended data Fig 5.**
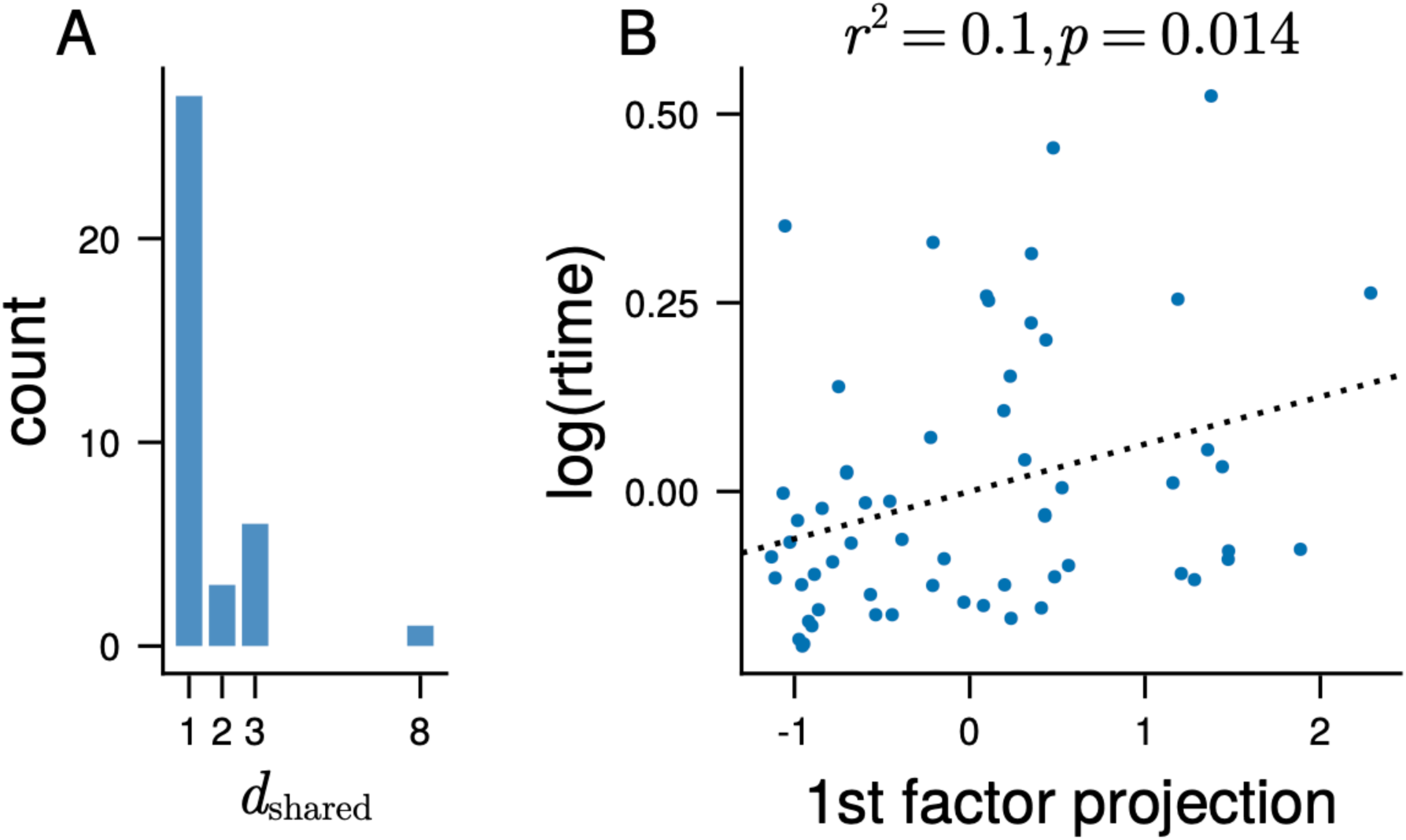
Factor analysis. **(A)** The distribution of the number of dimensions necessary to explain 95% of the shared variance in the post-go-cue period (0 to 50ms relative to go-cue onset) within each session and location. This number was determined via factor analysis, as in Khanna et al (2019)^18^. **(B)** The projection of population activity in the period from 0 to 50 ms relative to go-cue-onset for one session and one location onto the first factor component plotted against the logarithm of the reaction time. The projection exhibited a significant linear relationship with reaction time (r^2^ = 0.1, p=0.014).

## Notes

### Competing Interest Statement

The authors have declared no competing interest.

